# Effect of shear and tensile loading on fibrin molecular structure revealed by coherent Raman microscopy

**DOI:** 10.1101/2020.07.19.205005

**Authors:** Yujen Wang, Sachin Kumar, Arsalan Nisar, Mischa Bonn, Manuel K. Rausch, Sapun H. Parekh

## Abstract

Blood clots are essential biomaterials that prevent blood loss and provide a temporary scaffold for tissue repair. In their function, these materials must be capable of resisting mechanical forces from hemodynamic shear and contractile tension without rupture. Fibrin networks, the primary load-bearing element in blood clots, have unique nonlinear mechanical properties resulting from their hierarchical structure, which provides multiscale load bearing from fiber deformation to protein unfolding. Here, we study the fiber and molecular scale response of fibrin under shear and tensile loads *in situ* using a combination of fluorescence and vibrational (molecular) microscopy. Imaging protein fiber orientation and molecular vibrations, we find that fiber orientation and molecular changes in fibrin appear at much larger strains under shear compared to uniaxial tension. Orientation levels reached at 150% shear strain were reached already at 60% tensile strain, and molecular unfolding of fibrin was only seen at shear strains above 300%, whereas fibrin unfolding began already at 20% tensile strain. Moreover, shear deformation caused progressive changes in vibrational modes consistent with increased protofibril and fiber packing that were already present even at very low tensile deformation. Together with a bioinformatic analysis of the fibrinogen primary structure, we propose a scheme for the molecular response of fibrin from low to high deformation, which may relate to the teleological origin of its resistance to shear and tensile forces.

**Graphical Abstract:** 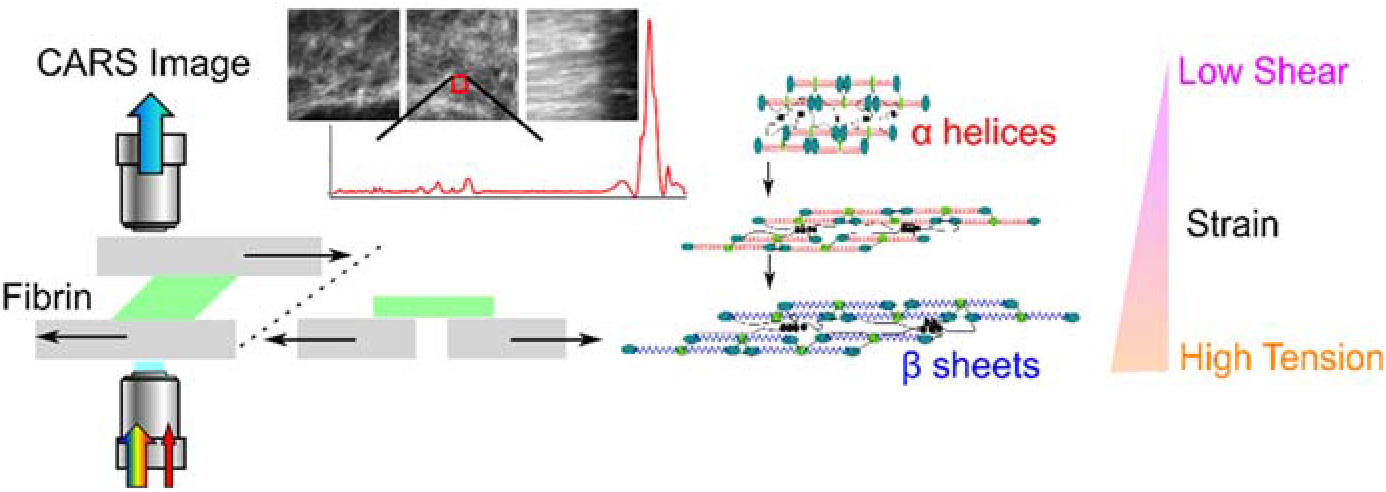

## Introduction

Fibrin is the three-dimensional (3D) mesh-like, protein network that comprises the bulk mass of blood clots and gives clots their mechanical strength [1,2]. Fibrin networks are formed by polymerization of fibrin molecules that are produced when monomeric (soluble) fibrinogen is enzymatically cleaved by thrombin into fibrin during the clotting cascade [3,4]. This cleavage initiates polymerization of fibrinogen monomers into a hierarchical 3D fibrin network to prevent blood loss at the wound site [5–7]. In addition to preventing blood loss by forming a physical barrier, fibrin also provides mechanical stability to blood clots. Fibrin clots must be able to sustain mechanical forces such as muscle and wound contraction (tension) and blood flow (shear) from various directions during wound healing to avoid additional hemorrhaging or thrombus formation [8–10].

Blood clots, as well as fibrin networks (or hydrogels), have been studied extensively using mechanical testing [7–9,11–20]. Similar to other biopolymers such as actin, collagen, and vimentin, fibrin networks and whole blood clots exhibit elastic properties that show strain stiffening behavior under increasing shear [7,19,21,22] and tensile [15–17,23,24] loading, meaning that fibrin biomaterials show an increasing elastic modulus with deformation. Such nonlinear elasticity is linked to hemostasis, blood clot stability, and supportive function for wound healing under dynamic loading environments [7,8,19,25]. As fibrin is a hierarchical material, with monomers assembling into protofibrils, protofibrils bundling into fibers, and fibers entangling into the 3D meshwork, the biomechanical properties of fibrin are interconnected from the molecular level to the fiber level [7,10,26–28]. Therefore, understanding the mechanics of fibrin networks at the molecular level can provide fundamental knowledge about the biomechanics of blood clot stability.

Compared to the number of studies relating macroscopic material properties with changes in fibrin at the fiber level (easily in the hundreds), e.g. fiber diameter, branching, or mesh porosity, relatively few studies have focused on molecular-scale changes in fibrin as a function of deformation. In both simulation and experimental studies, fibrin molecules have been shown to unfold under tensile force with the alpha-helical coiled-coil region contributing the primary resistance to deformation [15–18,24,29]. However, other experimental work has shown that unstructured αC domains were directly involved in load-bearing with the coiled-coil regions playing a minimal role [28,30–33]. Unfortunately, these studies were executed in very different experimental geometries, and the loading regimes are not fully comparable, resulting in an unresolved debate about how fibrin accommodates mechanical loads. Therefore, a study comparing the molecular scale response of fibrin from small to large deformations in the same experimental system is needed to help clarify the molecular response of fibrin to mechanical loads.

Experimental tools used to study the molecular response of fibrin include atomic force microscopy (AFM), scanning electron microscopy (SEM), and spectroscopic techniques. AFM studies have provided force-extension measurements capable of showing protein unfolding on fibrinogen monomers *in situ*. Measurements on oligomers, even up to single fibrin fibers [17,34,35], measure very fine mechanics but no longer offer the molecular-level insights of single-molecule experiments. In SEM, one can visualize single fibers – also under load [36], but originally hydrated fibrin networks are most often dehydrated and fixed, and combining SEM with *in situ* mechanical loading is not trivial. Spectroscopic tools such as Fourier transform infrared (FTIR) and Raman scattering offer alternative methods to probe structural changes of proteins using molecular vibrations. These methods can be performed on 3D fibrin hydrogels under mechanical deformation *in situ*. Litvinov *et al*. analyzed the Amide I and Amide III vibrational bands of blood clots under large extensional strains from attenuated total reflectance FTIR (ATR-FTIR), revealing the coiled-coil region of fibrin unfolded into β-sheets under large tensile deformation [18]. Similarly, Fleissner *et al.* used coherent Raman imaging to show the *in situ* spatial heterogeneity of α-helices and β-sheets in tension-stretched fibrin composite hydrogels [23]. Both Litvinov *et al.* and Fleissner *et al.* used tensile (or compressive) loads and only considered the amide spectral region to resolve α-helix to β-sheet unfolding of fibrin molecules.

In this study, we used broadband coherent anti-Stokes Raman scattering (BCARS) microscopy and a custom-built loading device to image the molecular vibrational signature of fibrin under shear and tensile deformations *in situ*. We analyzed the hyperspectral datasets in the C-H stretch (2800 – 3000 cm^−1^), phenyl ring (1000 – 1050 cm^−1^), and Amide I (1630 – 1700 cm^−1^) regions. Our results show that shear loading resulted in distinct molecular changes in fibrin consistent with increasing molecular packing and minimal protein unfolding, which was only observed above 300% strain. On the other hand, β-sheet signatures from fibrin unfolding were observed in tension already at 20% strain. Fluorescence images of deformed fibrin showed that shear deformation caused substantially less fiber alignment compared to tensile deformation at similar strains. Our results suggest a simultaneous fiber alignment and molecular packing in fibrin at lower forces with fibrin unfolding occurring at higher forces.

## Materials and Methods

### Fibrin network preparation

Fibrin networks were prepared by dissolving fibrinogen (FB I, Enzyme Research Laboratories) in a buffer solution containing 20 mM HEPES and 150 mM NaCl at pH 7.4 to make a 20 mg/ml stock. The fibrinogen solution was mixed with thrombin in the buffer mentioned above to a final concentration of 1 U/ml, including 5mM CaCl_2_ (Human Alpha Thrombin, Enzyme Research Laboratories) to obtain a pre-gel fibrin solution. The fibrin concentrations were 5 mg/ml for confocal microscopy and 15 mg/ml for BCARS (to increase signal strength).

For BCARS microscopy of sheared samples, the pre-gel fibrin solution was placed at the center of a detergent (Micro-90, Sigma)-cleaned glass slide with a 10 mm circle marked using a hydrophobic marker to prevent the pre-gel fibrin solution spreading. Three coverslips (20 × 20 mm^2^ with a total thickness of ~ 0.45 mm) were placed at the two sides of a glass slide and used as spacers for the network. The pre-gel solution was then covered with a cleaned larger coverslip (24 × 60 mm^2^, ~ 0.15mm thick). The entire sandwich was incubated in an oven at 37°C for two hours to gel. After two hours, two coverslip spacers on each side were carefully removed without disturbing the network to accommodate water evaporation. The sample was further allowed to gel at 37°C for an additional 4-6 hours. The final fibrin network had dimensions of ~ 10 mm diameter and ~ 130 µm height as determined by optical measurements (**Fig. S1)**. The entire preparation process is shown in **Figure S2A and S2B**.

Similarly, for BCARS microscopy of tensed samples, a pre-gel solution (also 15 mg/ml fibrinogen) was placed on a homemade Teflon plate with a 3 mm wide center ridge that accommodates a coverslip (#1.5, 24×36 mm^2^) on either side of the ridge such that the profile along the surface was flat. The pre-gel solution was allowed to spread within a hydrophobic pen-marked rectangular region (10 mm × 20 mm in width and length, **Fig. S2C**) and incubated at 37°C for gelation. After two hours, the coverslip-network-coverslip set was carefully removed from the Teflon plate to minimize sample perturbation, and a larger coverslip (#1.5 24 × 60 mm^2^) was used to form a chamber around the bottom of the network. Two steel spacers (3 × 15 × 0.5 mm^3^) were placed beside the network to support another coverslip on top that provided a flat surface for transmitted CARS imaging (**Fig. S2D**).

### Microscopy

#### Confocal fluorescence microscopy

Fibrin morphology (5 mg / mL) under shear and tensile deformation was imaged using a laser scanning confocal microscope (FV3000, Olympus) with a 60X, 1.1 NA (LUMFLN60XW, Olympus) water dipping objective lens. Fibrin networks were made with 9:1 ratio of unlabeled fibrinogen:Alexa 488 labeled fibrinogen (Sigma Aldrich). For shear testing, a home-built deformation stage was used during experiments (**Fig. S3A**), and Z-stacks with 0.5 µm step size were acquired at least 200 µm from the glass surface into the network. The pinhole on the microscope was always set to 1 Airy unit. Each acquired stack was processed by ImageJ (using the “Image reslice” and “Group Z projection commands with “maximum intensity” option), producing a set of XZ images. Both XY and XZ images were further analyzed in OrientationJ [37,38] for fiber orientation with a 5º bin size. We note that for linear shear, the loading axis changes with shear strains in the XZ plane [39]. For tension, the same setup was used but in a different configuration (**Fig. S3B**). Z-stacks were not acquired in tension.

For thioflavin T (ThT, Sigma) experiments, the dye was prepared at 100 µM according to supplier’s instructions. The ThT solution was added and incubated for 30 min. Afterward, the ThT solution was wicked away with Kimwipes, and the sample was washed several times by the same HEPES buffer mentioned in the fibrin preparation section above. Samples were excited at 488 nm, and emission was collected from 530 to 580 nm, separate from fluorescent fibrinogen. Images were processed by ImageJ to subtract background fluorescence.

#### BCARS hyperspectral microscopy

For molecular microscopy, we used the BCARS hyperspectral microscope that was previously used for fibrin measurements [23]. Briefly, an Nd:YAG microchip laser producing nanosecond pulses (~ 30 KHz repetition rate) at 1064 nm and a broadband supercontinuum with a spectrum ranging from 1100 nm to 2400 nm (Leukos CARS, Leukos) was used for excitation. The two beams were focused by either a 100X, 0.85 NA objective (LCPLN100XIR, Olympus) for shear or a custom-built 60X, 1.1 NA objective (Special Optics) for tension. Different objectives were used due to the different working distances required for the two setups. The signal was collected in transmission by a 20X, 0.4 NA objective (M-20X, MKS Newport). The sample was moved to achieve a pixel size of 0.5 × 0.5 um^2^, and the CCD integration time for each spectrum with ~ 400 ms integration time maximize signal while avoiding saturation.

For shear measurements, the fibrin construct (shown in **Fig. S2B**) was carefully mounted on the BCARS microscope (See **Fig. S3C**). The bottom glass slide was fixed, and the top coverslip was mounted to an electronic screw (Z825B, Thorlabs). The speed and acceleration were set as 0.2 cm/s and 0.1cm/s^2^, respectively. Shear was applied by moving the top coverslip by *Δx* relative to the fixed glass slide and the strain was calculated as 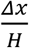 where *H* was the height of fibrin network. *H* was measured using the microscope, assuming the refractive index of the fibrin network was 1.33 (the same for water) as described in **Figure S1 and Equation S1, S2**. We performed BCARS measurements near the middle of the fibrin network to avoid edge effects but report strain using the equation above for simplicity.

For tension BCARS measurements, the fibrin network was mounted on the BCARS microscope, as shown in **Figure S3B**. One end of the network construct was fixed while the other end was connected to the electronic micrometer. Uniaxial tension on the network was applied by moving the micrometer (ΔL), and the strain was calculated as ΔL/L where L was the initial gauge length of the fibrin network. During both shear and tension experiments, a spectral signal was collected using BCARS from the bulk network (at least 50 µm from the glass surfaces) to avoid any unexpected effects from interfacial fibrin on the glass surfaces.

### CARS Data Processing

BCARS raw spectra were converted into Raman-like spectra for quantitative analysis, as previously reported in literature [40,41]. BCARS spectra were collected over a range from 700 cm^−1^ to 4000 cm^−1^ with an average spectral resolution of 4 cm^−1^. After phase retrieval of the Raman line shape, all spectra were normalized by the most intense vibration of the CH_3_ vibration (2935 cm^−1^ for tension, and 2970 cm^−1^ for shear) in order to account for scattering by the sample. For each loading condition, an average spectrum was calculated by averaging spectra from 81 × 81 spatial positions in each hyperspectral image. From the spectral data, three analytical quantities were used for molecular analysis of fibrin: 1) ratio of the CH_3_ symmetric (2935 cm^−1^):asymmetric (2970 cm^−1^) stretch [42–47]; 2) ratio of the Amide I α-helix (1649 cm^−1^):β-sheets (1672 cm^−1^) [48]; and 3) the ratio of two phenylalanine peaks (1004 cm^−1^ and 1045cm^−1^) [49]. The peak widths for integrating the two CH_3_ stretching peaks were 24 cm^−1^ around the center wavenumbers. For the ratio of α-helix to β-sheets, the ranges for the two peaks were 1645 cm^−1^ to 1655 cm^−1^ and 1669 cm^−1^ to 1679 cm^−1^, respectively. The two phenylalanine peaks were not equal in their peak widths, so the widths were 30 cm^−1^ and 54 cm^−1^ for 1004 cm^−1^ peak and 1045 cm^−1^ peak, respectively (See **Equation S3, S4 and S5** for ratio details).

## Results

### Shear and tensile strains induce alignment of fibrin fibers at different strains

We initially observed fibrin morphology changes under mechanical deformation – both shear and tension – using confocal fluorescence microscopy. Freshly prepared fibrin networks showed a porous network, and fibers were distributed isotropically in both the XY and XZ planes (**Fig. 1A and 1B, 0%**). During shear deformation, fibrin fiber alignment was detected more strongly axially (in XZ) compared to laterally (in XY) at lower strain. Quantitatively, the histogram of fiber alignment in the XZ plane at 50% shear showed close to 18% alignment, almost triple that of the 6% alignment in the XY plane (**Fig. 1C and 1D**). The maximum alignment in the XY plane reached 14% at 250% strain (above which this network detached at higher strains) while 30% alignment was detected in the XZ plane for the same shear strain. Conceptually, a viscoelastic network under simple shear in the horizontal direction with a defined gap will tend to build stress on fibers in the XZ plane. Semiflexible fibers like fibrin rotate according to the stress direction, which drives the alignment of the fibers along the loading direction in the XZ plane. Such behavior has been shown to be quantitatively accurate for predicting elasticity and network behavior of semiflexible actin filaments [50].

**Figure 1.**
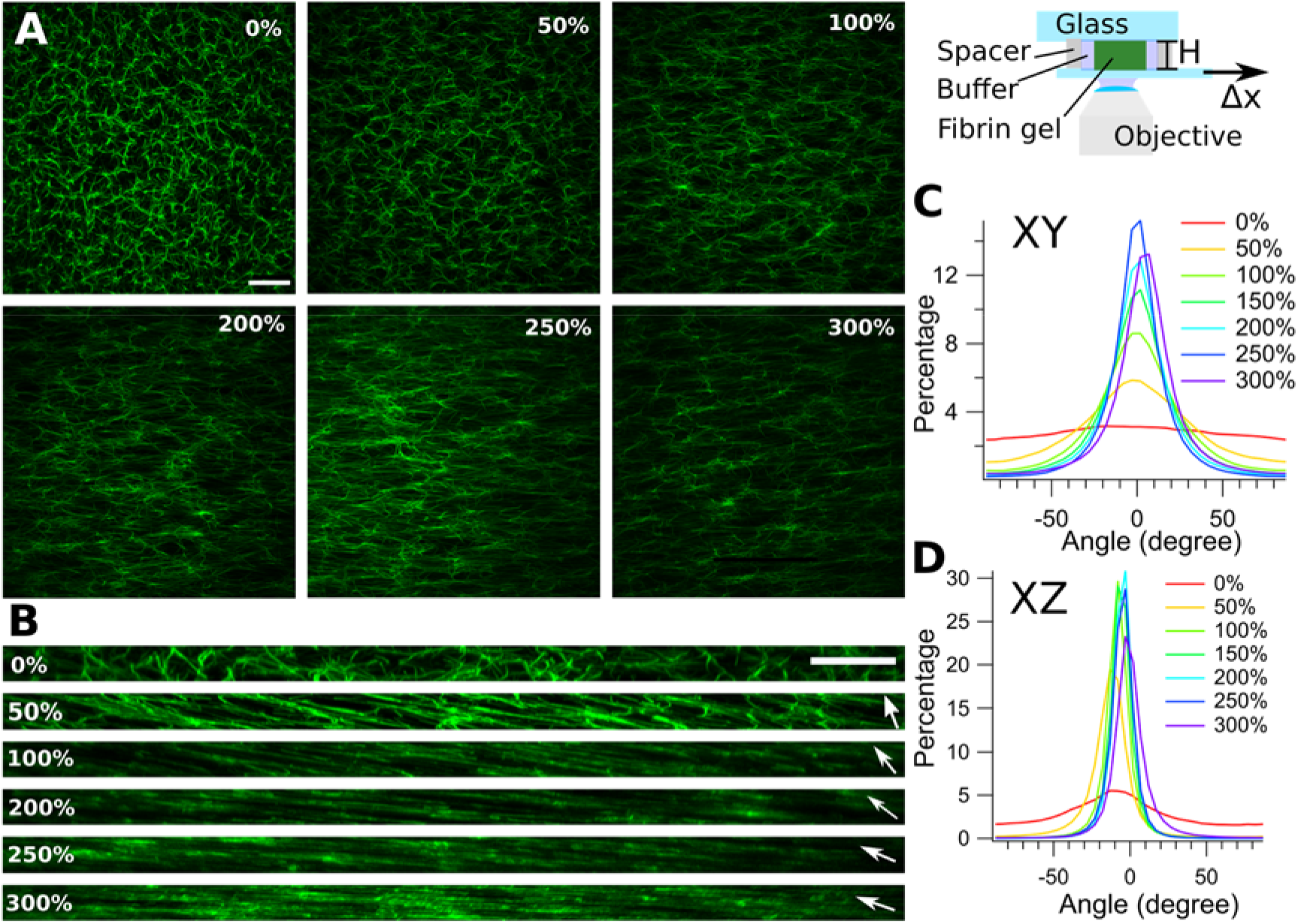
Confocal fluorescence microscopy of fibrin network under simple shear. Top right image shows the experimental geometry. **(A)** XY confocal images of fibrin doped with fluorescent fibrinogen under shear deformation. Each image is maximum intensity projected over 10 μm. **(B)** XZ images of resliced, projected confocal images of fibrin reconstructed from Z stacks of XY images. White arrows show the loading axis projected into the XZ plane. **(C)** Histogram of XY fiber orientation angles at increasing shear strain. **(D)** Histogram of XZ fiber orientation angles at increasing shear strain. The y-axis for the histograms corresponds to the percentage of fibers out of the total number of fibers detected binned in 5^◦^ steps. Scale bars are 30 μm (A) and 20 μm (B).

Under tension, fibrin fibers showed a classic uniaxial tensile behavior, with elongation in the (horizontal) loading direction clearly visible in the XY planes (**Fig. 2A**). Randomly oriented fibrin fibers started to orient in the stretching direction at strains as low as 20%; alignment was visible in XY images as well as in the orientation histogram, where a strong peak near 0^◦^ showed ~ 10% fiber alignment (**Fig. 2B)**. Further tensile strain resulted in an increased alignment of fibrin fibers in the stretch direction and a narrowing of the orientation angular distribution. In addition, fibers started to pack closer and align nearly parallel to each other at 60% tensile strain, where the XY alignment reached almost 14%. Tensed fibrin samples often fractured above 100% tensile strain. These confocal microscopy results show that fibrin under shear and tensile strain aligned in the loading direction, as expected, with tensile loading showing earlier alignment – at lower strains – in comparison to shear.

**Figure 2.**
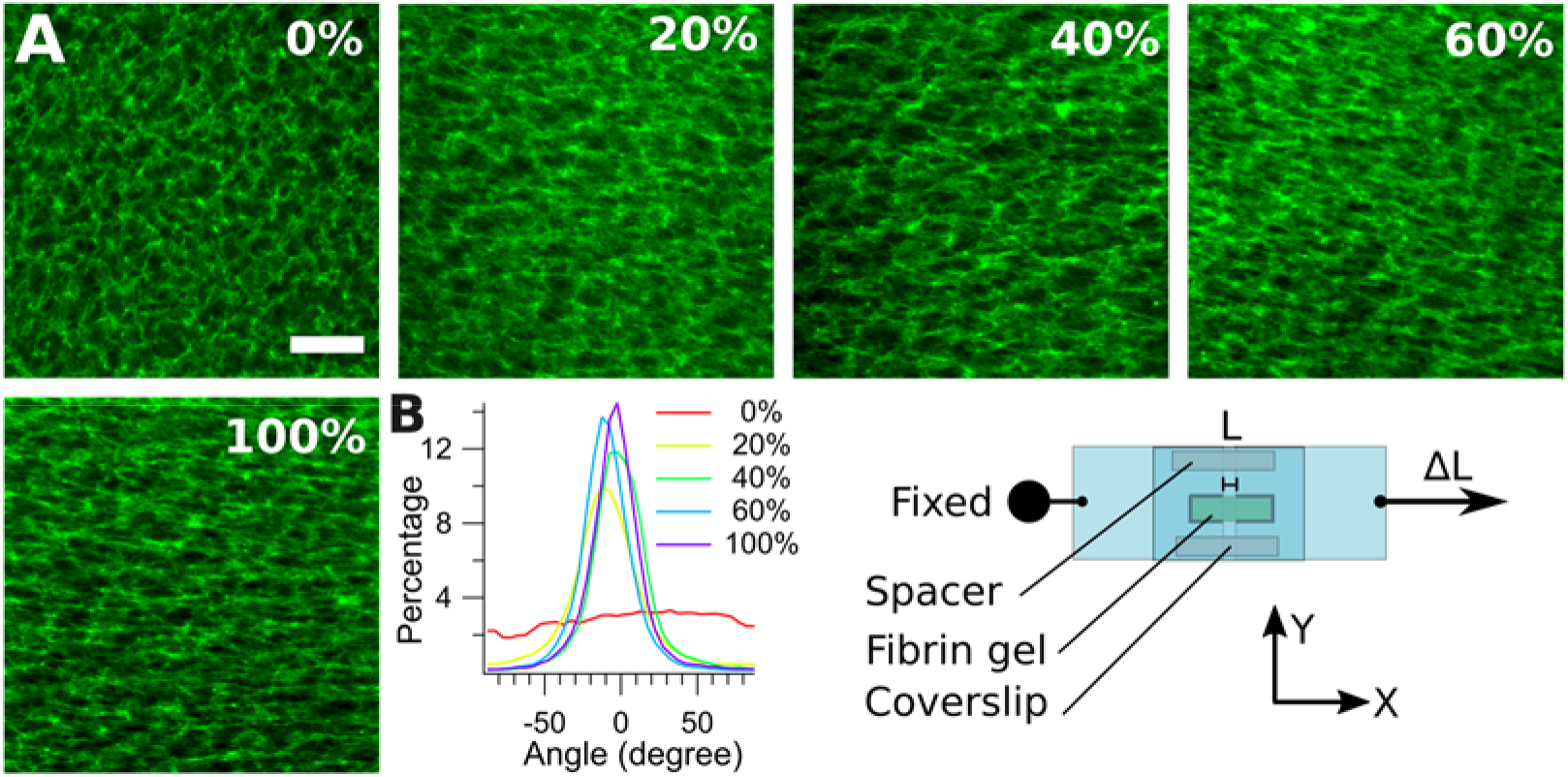
Confocal microscopy of fibrin networks under uniaxial tension. Bottom right image shows the experimental geometry. **(A)** Single confocal images of fibrin doped with fluorescent fibrinogen under increasing tensile strain: 0%, 20%, 40%, 60% and 100%. Scale bar is 20 μm. **(B)** Histogram of fiber orientation (with respect to horizontal/loading direction) for images shown in A. The y-axis is the percentage of fibers out of the total number of fibers detected binned in 5^◦^ steps.

### Shear strain causes increased packing of CH_3_ groups in fibrin molecules

Following confocal microscopy, we collected information on the molecular changes in fibrin under shear loads using BCARS vibrational microscopy with an *in situ* loading device as described in the methods (see **Fig. S3A**). BCARS hyperspectral data cubes, consisting of a vibrational spectrum at each spatial position, were collected from fibrin networks prepared similarly to those used for confocal microscopy. For shear measurements, we acquired multiple (N > 5) hyperspectral datasets in increments of 50% shear strain after waiting 10 minutes upon reaching the desired strain level, and a representative dataset is shown here. The resulting BCARS spectra, after retrieval of the Raman features, were analyzed by plotting the histogram of respective peak ratios to quantify the molecular heterogeneity and changes with increasing shear strain. As a starting point, we mapped the CH_3_ stretching ratio (symmetric CH_3_ / asymmetric CH_3_) as a function of shear strain, as shown in **Figure 3A**. Unstrained fibrin showed a mean CH_3_ stretching ratio of 0.75 that increased to 0.85 at medium strain (150%) and later reached ~ 1 for highly (400%) sheared fibrin. The increased fibrin CH_3_ stretching ratio with shear strain shows that upon shearing, fibrin intensity for the symmetric CH_3_ stretching increased, whereas asymmetric CH_3_ stretching mode decreased. Average spectra from hyperspectral datasets (81 × 81 pixels) with gray boxes show the three regions of interests: phenylalanine peaks, the Amide I region, and the C-H stretch (**Fig. 3B**). Ratio histograms (**Fig. 3C, 3D, 3E**) were plotted, and parameters from Gaussian fits to all histograms are shown in **Table S1**.

**Figure 3.**
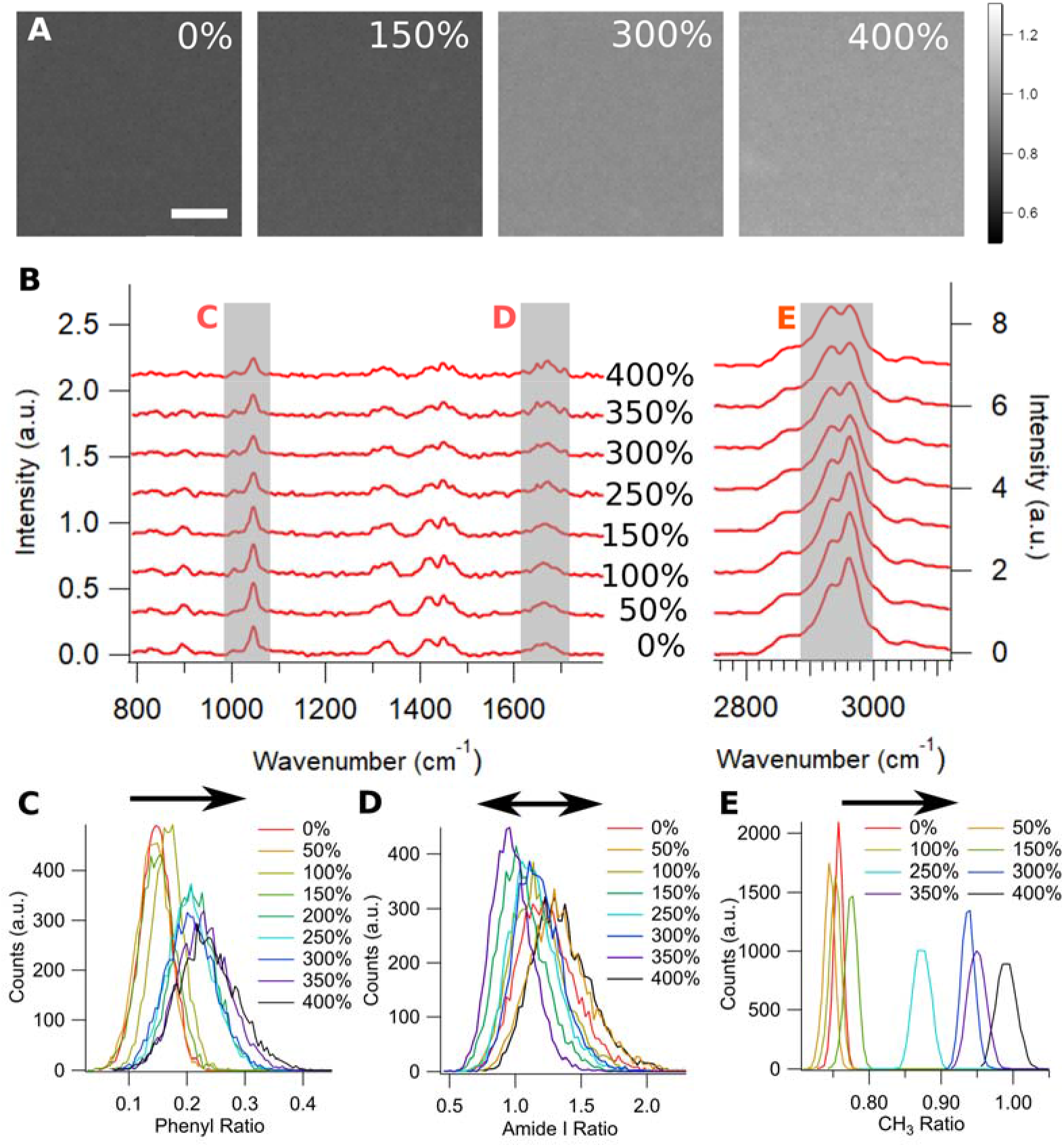
BCARS molecular spectroscopic imaging of fibrin under simple shear. **(A)** A representative set of CH_3_ ratio images of fibrin; intensity values are from Isymmetric CH_3_ / Iasymmetric CH_3_). The images were 40 × 40 μm^2^ with 81 pixels along both x- and y-axis. The same color bar (far right) was used for all CH_3_ ratio images. Percentages show the amount of shear strain. Scale bar is 10 μm. **(B)** Complete spectral information obtained by averaging 81 × 81 pixels from hyperspectral images of a fibrin network under different shear strain. Red letters correspond to panels C, D, and E below. **(C, D and E)** Histograms showing of phenylalanine peak, Amide I, and CH_3_ ratios, respectively. Each loading group consisted of 6561 spectra for ratio calculations, and the bin sizes were 0.01. The arrows above the histograms show the trend of each histogram with increasing strain. Representative data is shown from N > 5 experiments.

The CH_3_ stretching modes originate from side chains of alkyl amino acids in proteins. The CH_3_ symmetric stretching (2935 cm^−1^) refers to all three C-H bond lengths vibrating in phase while the asymmetric stretching (2970 cm^−1^) C-H bonds length change in different phases. In fibrin under shear deformation, the CH_3_ asymmetric stretching was dominant at low strain, and the CH_3_ symmetric stretching mode increased while the asymmetric stretching mode decreased with strain, which shows the molecular environment in highly sheared fibrin is very different compared to unstrained fibrin. Histograms (**Fig. 3E**) of the CH_3_ ratio (symmetric CH_3_ / asymmetric CH_3_) from each pixel in the images show a clear trend of increasing CH_3_ ratio and increasing width of these histograms with increasing shear strain. Based on confocal XZ images in **Figure 1B** showing that the network density increased at higher shear strain and established the connection between molecular packing and C-H stretch resonances, we surmise that the fibrin network became more tightly packed with increasing shear strain.

Similar to the CH_3_ stretching ratio, the phenylalanine peak ratio also changed with increasing shear strain. The phenylalanine peaks (at 1004 cm^−1^ and 1045 cm^−1^) represent two complex vibrational modes: the former is the symmetric ring breathing mode, and the latter is the ring C-H wagging mode [49,51]. These two phenyl modes can change with the molecular environment, indicating how the phenyl rings on amino acid side chains stack during deformation. The average spectra (**Fig. 3B**) show a slight increase in the phenylalanine peak at 1004 cm^−1^, while the peak intensity at 1045 cm^−1^ decreased more noticeably, resulting in an overall increase in phenylalanine peak ratio. The histograms for the phenylalanine peak ratio shifted to higher values with increasing strain, similar to that of the CH_3_ stretching ratio. Above 200% strain, the fibrin phenylalanine peak ratio started to broaden and increased substantially (**Table S1**, **Fig. 3C**).

The Amide I peak is a well-known indicator of protein secondary structure [47]. It is a vibrational combination of C=O and N-H in amino acid [52], which is sensitive to α-helices, unstructured loops, and β-sheets, with the peak locations at 1649, 1660 and 1672 cm^−1^, respectively [47]. However, the signal to noise ratio of Amide I region was low throughout the whole set of spectra such that multi-peak fitting was not stable. Therefore, we integrated the α helix (1643 – 1653 cm^−1^) and β-sheet (1669 – 1679 cm^−1^) portions of the spectra and took their ratio, which has been previously used to quantify changes in protein secondary structures [18,23,52]. Compared to the other two ratios, the Amide I ratio did not display a clear trend, both increasing and decreasing as the strain on the network was increased. The ratio increased with shear from 0% to 50%, 150% to 250%, 250% to 300%, and 350% to 400% but decreased from 50% to 100%, 100% to 150% and 300% to 350%. As for the width, the full with at half maximum (FWHM) slowly decreased until 350% and increased again at 400%, possibly due to the network detaching from the plates (see **Table S1**). We attempted to look for consistent trends in the Amide I ratio by grouping the Amide I ratios into three strain categories: low strain (0% to 50%), medium strain (100% to 200%), and high strain (250 – 350%, discarding 400%). For low and medium strain, the average of the Amide I ratio was 1.16 ± 0.08 (mean ± std. dev), and the FWHM was 0.400 ± 0.031 (mean ± std. dev). For the high strain group, we detected a slightly decreased and narrower Amide I ratio, where the average ratio was 1.07 ± 0.1 (mean ± std. dev) and the FWHM was 0.385 ± 0.028 (mean ± std. dev), respectively. No statistically significant differences were observed for the Amide I ratio values between the low, medium and high strain levels. This suggests that even at high shear strain (up to 350%), only very limited changes in the secondary structure were observed.

### Fibrin unfolds under tension but is unaccompanied by other molecular changes seen in shear

We performed similar BCARS hyperspectral imaging experiments for fibrin under uniaxial tension. **Figure 4A** shows images of the integrated C-H stretch (2800 cm^−1^ – 3000 cm^−1^) intensity in fibrin from BCARS hyperspectral data of 0% nominal strain to 100% strain. tensed fibrin. Here, strain values refer to nominal strain; a small amount of strain applied to the fibrin samples during mounting was inevitable. Fibrin at 0% nominal strain showed a network of fibers oriented randomly, similar to the confocal images shown earlier. Similarly, we observed increasing fiber alignment along the horizontal (loading) direction with increasing strain. At 100% nominal strain, we observed very strong alignment and increased fiber density in the network.

**Figure 4.**
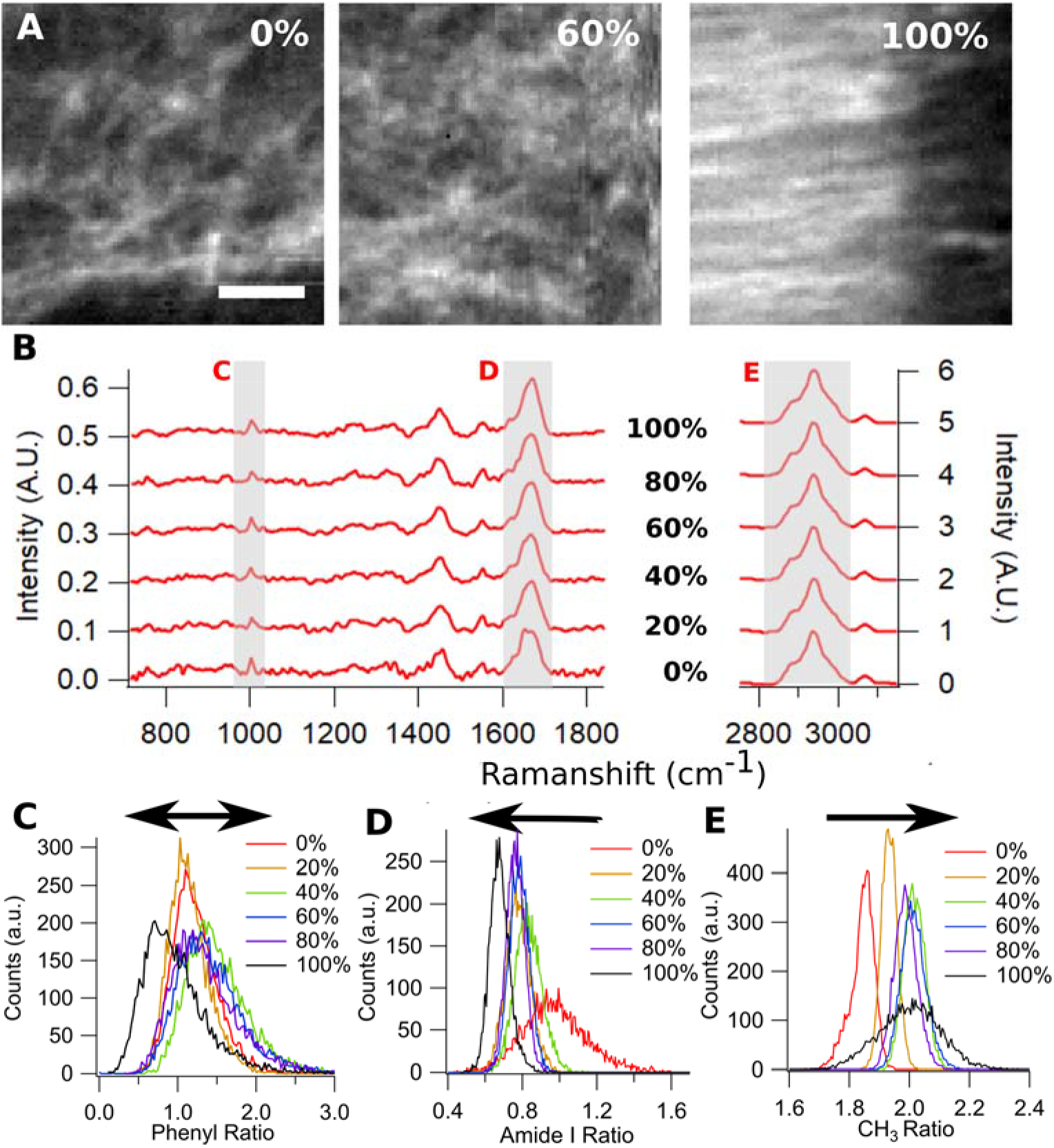
BCARS molecular spectroscopic imaging of fibrin under uniaxial tension. **(A)** Fibrin CH images from BCARS data (values are integrated intensity from 2800 cm^−1^ – 3000 cm^−1^) from 40 × 40 μm^2^ with 81 pixels along both x- and y-axis. Strain was applied along the horizontal direction. Percentages show the amount of nominal tensile strain. Scale bar is 10 μm. **(B)** Complete spectral information obtained by averaging 81 × 81 pixels data points from hyperspectral images of a fibrin network under increasing tension. Red letters correspond to panels C, D, and E below. (**C, D and E)** Histograms showing the phenylalanine, Amide I, and CH_3_ ratio, respectively. Each group consisted of 6561 spectra for ratio calculation, and the bin sizes were 0.025, 0.01, and 0.01 for phenylalanine ratio, Amide I, and CH_3_ ratio, respectively. The arrows above the histograms show the trend of each histogram with increasing strain. Representative data is shown from N > 5 experiments.

The average spectra for fibrin at increasing tensile strains (**Fig. 4B**) shows that fibrin networks under increasing tensile deformation exhibited minimal changes in their molecular signature outside of the Amide I band. The CH_3_ and phenylalanine peaks appeared almost constant with increasing strain. The CH_3_ symmetric stretching mode (2935 cm^−1^) was very strong, and the asymmetric stretching mode (2970 cm^−1^) was only barely visible as a shoulder beside it, even at 0% nominal strain, which shows that sample handling in tension experiments already caused changes in network structure that are on par with very large shear strains. The average CH_3_ ratio was ~ 2 at 100% nominal tensile strain, which was 2-fold higher than the in shear at 350% strain. The histograms of the CH_3_ ratio from tensile-loaded fibrin show that the average CH_3_ ratio value increased from 1.8 to 2 with strain (**Fig. 4E**). Next, for phenylalanine ring modes, we only observed the 1004 cm^−1^ peak; the 1045 cm^−1^ peak was hidden within the noise, even at 0% nominal strain. Therefore, the phenylalanine ratio in tension was highly susceptible to noise; even so, the average values at all tensile deformations were ~ 2-fold more than for shear with no clear trend as a function of increasing tensile strain (**Fig. 4C**). Finally, we observed that the Amide I ratio (α-helix:β sheet) changed with increasing tensile strain, as expected. The intensities of Amide I were stronger compared to shear deformation, and the maximum of the Amide I band changed with strain, shifting from 1649 cm^−1^ to 1670 cm^−1^ (**Fig. 4B**), showing an increased presence of β sheets. This result is consistent with Litvinov *et al*. and Fleissner *et al*. [18,23,48]. At 100% nominal strain, fibrin fibers showed that the average Amide I ratio decreased by 35% compared to the unloaded case. Overall, an increase in tensile strain (0% - 100%) not only decreased the Amide I ratio histogram average value, but also resulted in a decreased Amide I ratio histogram width (**Fig. 4D**), suggesting that protein structure was more homogenous at high strain in fibrin networks under tensile force. In summary, the subtle molecular changes observed in the CH_3_ and phenylalanine peaks in sheared fibrin are hardly discernible under tension while changes in the secondary structure are clear in tension but were somewhat obscured in shear.

### Thioflavin T staining confirms secondary structural changes in measured in BCARS

As we noticed differential changes in the secondary structure of fibrin under shear and tensile strains in our BCARS data, specifically the appearance of β-sheet structure, we used the molecular rotor dye Thioflavin T (ThT) as an additional probe for the presence of β-sheets in fibrin. ThT is a small molecule known for its high affinity to amyloid structures, consisting of more than 50% β-sheet, and has been used extensively to identify β-sheet-rich amyloid fibrils [53–55]. **Figure 5A** shows ThT staining of the same fibrin network at 0% and 200% shear strain under identical excitation and detector settings from confocal microscopy. We observed a slight decrease in ThT signal under high shear compared to 0% shear, indicating no increased β-sheet structure at 200% shear strain. In tension, on the other hand, the ThT dye was clearly increased at 75% strain over 0% strain when the excitation and detector settings were the same for both images (**Fig. 5B)**. In addition, we observed that strained fibers aligned nearly perfectly (along the loading direction) in the ThT Image. The ThT and BCARS imaging results are consistent with one another.

**Figure 5.**
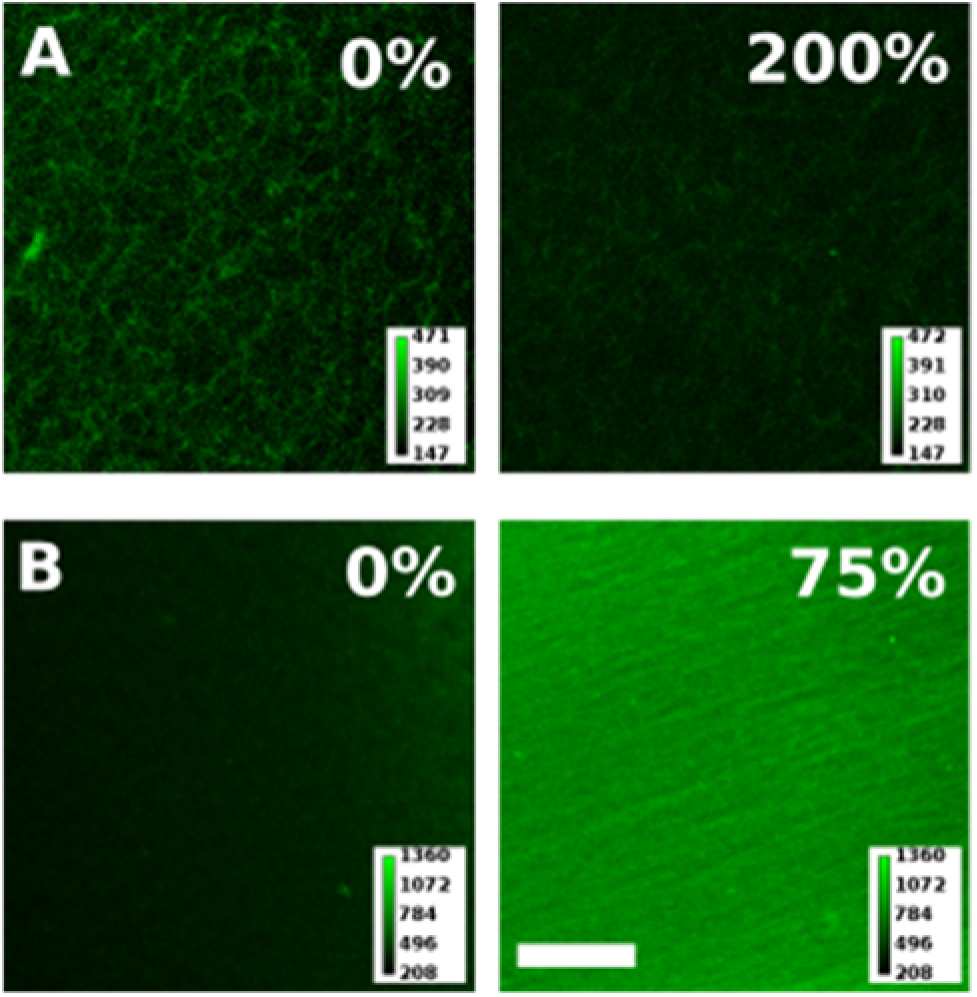
ThT fluorescence from fibrin under shear and tensile strain. **(A)** 0% and 200% shear strain were applied to the same fibrin network. The detector high voltage (HV) and excitation power were kept the same for both images. **(B)** 0% and 75 % tensile strain were applied to the same fibrin network. The detector HV and excitation power were the same for both images. Scale bar is 25 μm. The color scale (in counts) are shown in the bottom-right corner of each image.

## Discussion

### Analysis of amino-acid specific signals suggest fibrin molecular force accommodation mechanism

Vibrational microscopy of fibrin networks under shear and tensile deformation allowed us to probe molecular changes in fibrin from low to high forces. In the vibrational spectra, the CH_3_ stretching and phenylalanine ring modes showed clear changes with increasing shear strain. A logical follow-up question is: what is the origin of these signals? We trace these signals to the amino acids in fibrin from which they can arise. Phenylalanine is the only source for the phenyl ring mode peaks at 1004 cm^−1^ and 1045 cm^−1^; there are no other aromatic amino acid side chains in fibrin. The peak at 1004 cm^−1^ is the symmetric ring breathing mode, and the peak at 1045 cm^−1^ is the ring C-H wagging mode [49,51,56]. The ring breathing mode increased slightly with shear strain, while the ring wagging mode decreased more substantially. For the CH_3_ stretching, four amino acids in fibrin contain CH_3_ groups in their side chains: alanine (A) and methionine (M) has one CH_3_, and leucine (L), isoleucine (I), and valine (V) each have two CH_3_ groups. We located targeted amino acids in the fibrinogen #3GHG crystal structure [57–59] and highlighted the targeted amino acids in the protein model (**Fig. 6A and B**). We also counted the number of targeted amino acids according to the sequence for human fibrinogen on Uniprot [60–62], which is summarized in **Figure 6**.

**Figure 6.**
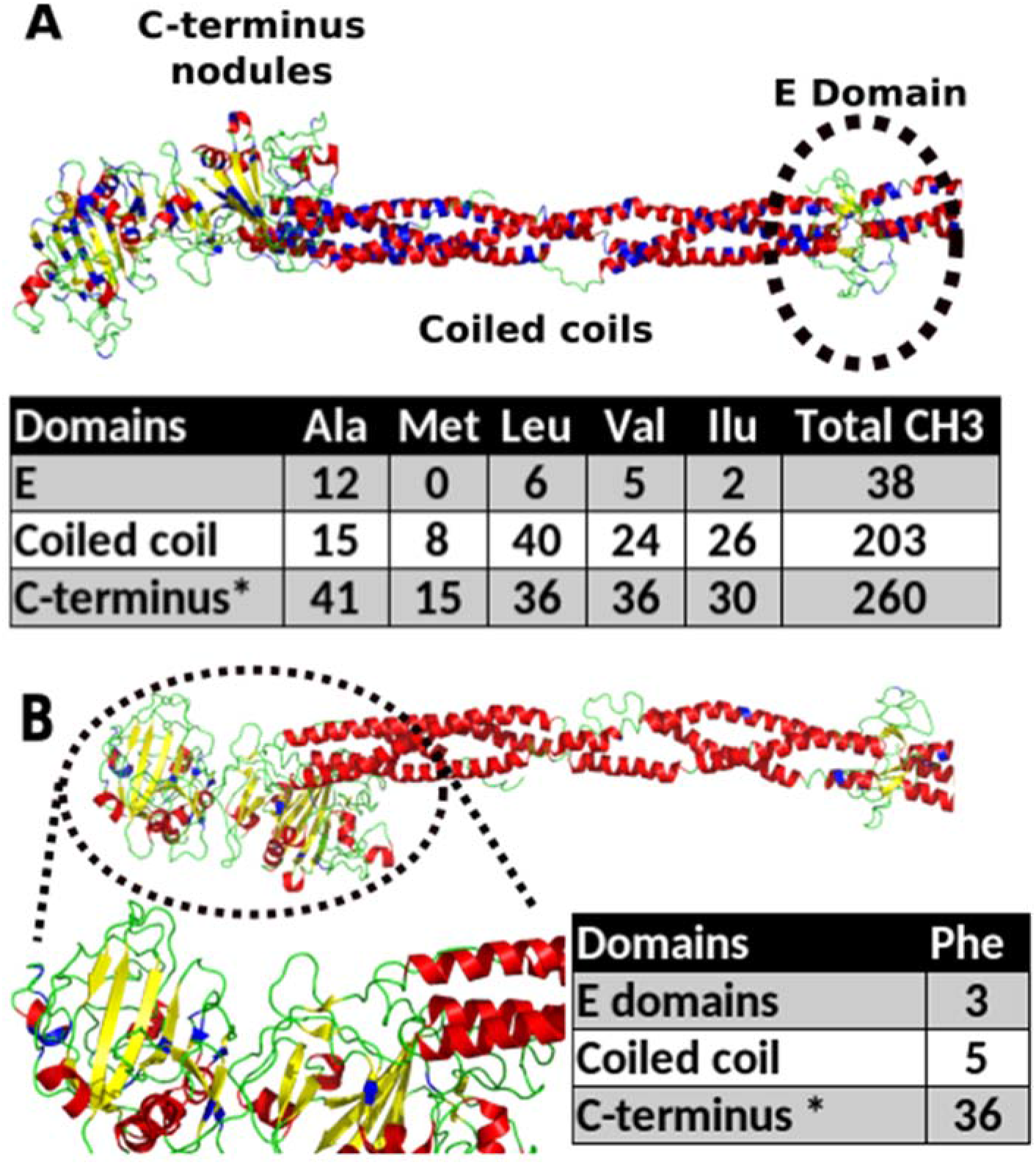
Targeted amino acid mapping on fibrinogen 3GHG model. (**A**) The distribution of amino acids with CH_3_ signal (Alanine, Valine, Leucine, Isoleucine) in fibrinogen. Red, yellow, and green colors correspond to α helixes, β sheets, and random coils, respectively. Methyl group-containing amino acids appear blue. We grouped the amino acids based on their presence in three different regions in fibrinogen: E domain, coiled coils, and C-terminal domains. **(B)** The distribution of phenylalanine in fibrinogen. The color logic was similar to (A), only in (B) phenylalanines were colored in blue. Most phenylalanine is in the C-terminal domains, so an enlarged picture is provided. * in the last row indicates that αC domains were counted even though they are not shown in the crystal structure.

In order to relate the spectral changes, especially those in shear, to potential changes in fibrinogen structure, we divided fibrinogen into three parts: E domains, coiled coils, and the C-terminal domains, which consists of β nodules, γ nodules, and αC domains. Interestingly, most of the phenylalanine is in β nodules, γ nodules, and αC domains, and only a few phenylalanines are located in coiled coil and E domain (**Fig. 6B, blue**). Therefore, when the phenylalanine peak intensities changed, we ascribe it as coming from the β nodules, γ nodules, and αC domains. Further support for this assumption comes from the observation that the phenylalanine peak ratio changes independently of the Amide I band, which shows that the coiled coil structure is not affected by shear strain, at least until very large strains. This suggests that smaller forces from shear deformation induce subtle conformational changes in the C-terminal domains as the phenylalanine ratio changes (**Fig. 6A**), consistent with the concept presented by Houser *et al.* where the αC domain was proposed to play a large role in load bearing[31].

A similar analysis of the CH_3_-containing amino acids (A, M, I, L, V) in fibrinogen molecule shows that the E domain, coiled coil, and C-terminal domains (including αC domain) contain 38, 203, and 260 CH_3_-containing amino acids, respectively (**Fig. 6A**). In vibrational spectroscopy, it is known that the local molecular environment strongly modifies C-H stretch modes [46,63,64]. C-H stretch modes have been used to indicate crystallinity and liquid/melt forms in various materials/chemicals, including n-alkanes [65], polyethylene [66], fatty acids [46] and lipids [67–70]. As these modes reflect changes over molecular distances, we speculate that the increasing CH_3_ ratio in shear comes from increased packing of fibers relative to another and increased packing of fibrin proteins in protofibrils within fibers.

As mentioned above, C-terminal domains are likely involved in the conformational rearrangements associated with phenylalanine ring mode changes during shear deformation. These domains also contain a large number of CH_3_-containing amino acids (209). Together with the observation that the Amide I region is reasonably constant until very large shear deformation (~300%), our results indicate that applying shear strain to fibrin mainly affects C terminal domains. In contrast, with minimal changes in the CH_3_ and phenylalanine ratios – but large changes in the Amide I ratio – under tension, where the force application more directly stretches fibers, we surmise that the coiled-coil region is primarily modified by tensile loading.

### Molecular changes from highly sheared fibrin are a precursor to secondary structure changes in tensed fibrin

**Figure 7A** shows a series of spectra from a fibrinogen solution, a native (unstrained) fibrin network, and fibrin networks with various levels of shear and tensile strain. We observed clear trends in the fibrin molecular response. The fibrinogen solution and native (unstrained) network appear very similar in all regions of the spectra. The spectra of fibrin under any deformation show larger amplitudes due to increased fibrin density in the network. Highly sheared fibrin and minimally tensed fibrin spectra also look very similar: the CH_3_ peak inversion compared to the fibrinogen or the unstrained network is clear, the presence of the 1004 cm^−1^ phenylalanine peak appears, and the Amide I shape is very similar. While these spectra show changes from additional vibrational modes, such as the Amide III (1260 – 1300 cm^−1^), methyl bending modes (1401 cm^−1^), CH_2_ deformation (1440 cm^−1^), and Amide II (1550 cm^−1^) [44–49], we focused here on the phenylalanine, CH_3_, and Amide I modes as the information content is duplicative for the other modes mentioned. Overall, the similarity between high shear and low tensile strain in CARS spectra shows that fibrin exhibits a similar molecular response under these conditions.

**Figure 7.**
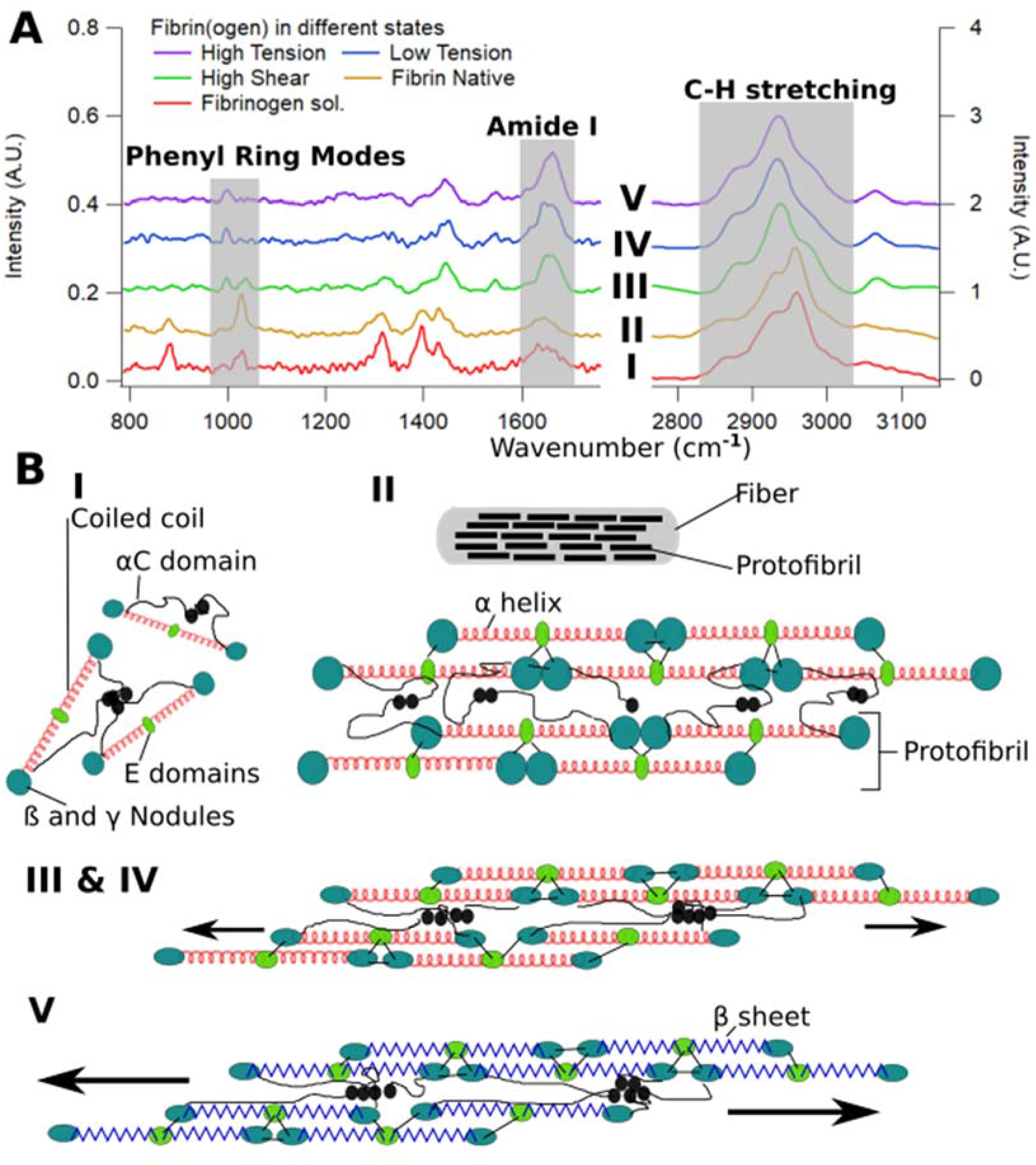
Summary of fibrin strain bearing mechanism. **(A)** BCARS spectra of Fibrin(ogen) in different states. Five spectra were shown from five different states of Fibrin(ogen): Fibrinogen solution (I), native (unstrained) fibrin network (II), highly sheared fibrin network (III), minimally tensed fibrin network (IV), and largely tensed fibrin network (V), shown from bottom to top. The three shaded regions show the C-H stretch, Amide I, and phenylalanine ring modes analyzed in this work. **(B)** Proposed mechanism for fibrin load bearing. The drawing shows states of strain matching with (**A**) by the same Roman numerals. (I) Soluble fibrinogen and (II) fibrin in the native (unstrained) state where the distances between fibers are large, and αC domains are loose. (III/IV) In the high shear strain / low tensile strain states, αC domains somewhat straighten and β and γ nodules deform. (V) In the highly tensed state, coiled coils in fibrin fibers unfold and undergo secondary structural transitions from α-helices to β-sheets. Parallel fibers were drawn to illustrate the concepts.

At high tension, the most noticeable change occurs for the Amide I shape. Therefore, it appears that the molecular changes seen during shear loading have already taken place for samples in our tensile experiments. We note that in shear experiments, the network was unperturbed prior to sample measurement; no additional sample handling or disruption was necessary. In tension experiments, we (gently) removed the network from the Teflon mold and mounted the network construct onto our loading setup on the microscope. This process, unfortunately, applied some irreversible loading to network, causing fibers to deform and possibly adhere to one another, as has been shown previously [19]. Therefore, the “0%” tension experiment is almost certainly not a “native” or an unstrained network, and this is potentially why spectra from 0% tension and 0% strain look so different.

Compared to shear loading, tensile deformation quickly aligns fibers in the XY plane – 20% tensile strain showed similar fiber alignment to 150% shear strain. Tensile deformation stretches fibers from end to end immediately, whereas the force axis rotates with increasing strain for simple shear. From the CARS shear data and analysis of the primary structure of fibrinogen, we deduce that changes in the phenylalanine peaks and the CH_3_ ratio likely originate from both nodules and αC region within the C-terminal domains as amino acids move/rearrange and fibers pack more densely.

The interpretation presented here is consistent with results from a previous study on fibrin structure under shear deformation (in the Couette geometry) using X-ray scattering by Vos *et al* [21]. In that study, the authors also found no clear evidence of secondary structural changes for shear deformation up to 300%. Moreover, they also deduced that the fiber density increased under shear partially mediated by the αC domain. In **Figure 7B**, we show a model of fibrin deformation, indicating molecular changes in fibrin with increasing deformation. Fibrinogen monomers and native fibrin networks show relaxed αC domains and comparatively larger distance between fibers and protofibrils (**Fig. 7B I, II**). At high shear/low tensile deformation, the network shows increased fiber/protofibril packing, β and γ nodules deformation, and αC domain deformation (**Fig. 7B III,IV**). Finally, under large tensile deformation, coiled coil regions unfold to accommodate additional forces (**Fig. 7B, V**). This mechanism is consistent with that predicted from molecular simulations of single molecule fibrin unfolding [16].

## Conclusion

In this work, we measured the molecular response of fibrin networks to simple shear and uniaxial tensile deformation with a combination of fluorescence and molecular microscopy *in situ*. Shear loading resulted in increased fiber density and molecular changes that reflect increased protein packing but very little change in secondary structure – up to 300% strain. On the other hand, the subtle molecular changes with increasing shear were nearly invisible in tension while pronounced changes fiber alignment, packing, and secondary structure were observed for tensile strains as low as 20%. The similarity of the high-shear and low-tension molecular responses suggest that forces applied to fibrin are initially accommodated by subtle rearrangement of fibrin C-terminal domains followed by unfolding of secondary structure in the coiled-coil domain under larger tensile loads. The results of this study are potentially helpful in the design of artificial wound dressing materials aimed at replicating the mechanical properties of fibrin clots.

## Supporting information

Supplementary Informatio

## Acknowledgements

We thank Florian Gericke and Marc-Jan van Zadel for technical support and Sabine Pütz for laboratory support. We also thank Dr. Frederik Fleissner and Dr. Jenée Cyran for critical reading of the manuscript and fruitful discussions. S.K. acknowledges Alexander von Humboldt Foundation Postdoctoral Fellowship, and S.H.P acknowledges support from the Welch Foundation (F-2008-20190330), and the Human Frontiers in Science Program (RGP0045/2018).

## References

[1] S.T. Lord, Fibrinogen and fibrin: scaffold proteins in hemostasis:, Current Opinion in Hematology. 14 (2007) 236–241. https://doi.org/10.1097/MOH.0b013e3280dce58c.

[2] A.S. Wolberg, Thrombin generation and fibrin clot structure, Blood Reviews. 21 (2007) 131–142. https://doi.org/10.1016/j.blre.2006.11.001.

[3] M.W. Mosesson, Fibrinogen and fibrin structure and functions, J Thromb Haemost. 3 (2005) 1894–1904. https://doi.org/10.1111/j.1538-7836.2005.01365.x.

[4] A. Undas, R.A.S. Ariëns, Fibrin Clot Structure and Function: A Role in the Pathophysiology of Arterial and Venous Thromboembolic Diseases, Arterioscler Thromb Vasc Biol. 31 (2011). https://doi.org/10.1161/ATVBAHA.111.230631.

[5] B. Blombaeck, B. Hessel, D. Hogg, L. Therkildsen, A two-step fibrinogen-fibrin transition in blood coagulation, Nature. 275 (1978) 501–505.

[6] S.J. Everse, G. Spraggon, L. Veerapandian, M. Riley, R.F. Doolittle, Crystal Structure of Fragment Double-D from Human Fibrin with Two Different Bound Ligands†, Biochemistry. 37 (1998) 8637–8642.

[7] I.K. Piechocka, R.G. Bacabac, M. Potters, F.C. MacKintosh, G.H. Koenderink, Structural Hierarchy Governs Fibrin Gel Mechanics, Biophysical Journal. 98 (2010) 2281–2289. https://doi.org/10.1016/j.bpj.2010.01.040.

[8] R.W. Rosser, W.W. Roberts, J.D. Ferry, RHEOLOGY OF FIBRLN CLOTS. IV. DARCY CONSTANTS AND FiBER THICKNESS, Biophysical Chemistry. 7 (1977) 153–157.

[9] C. Gerth, W.W. Roberts, J.D. Ferry, RHEOLOGY OF FIBRIQI CLOTS. II. LINEAR VlSCOELASTIC BEHAVIOR IN SHEAR CREEP, Biophysical Chemistry. 2 (1974) 208–217.

[10] J.W. Weisel, The mechanical properties of fibrin for basic scientists and clinicians, Biophysical Chemistry. 112 (2004) 267–276. https://doi.org/10.1016/j.bpc.2004.07.029.

[11] M.K. Rausch, J.D. Humphrey, A microstructurally inspired damage model for early venous thrombus, Journal of the Mechanical Behavior of Biomedical Materials. 55 (2016) 12–20. https://doi.org/10.1016/j.jmbbm.2015.10.006.

[12] P.A. Janmey, E.J. Amis, J.D. Ferry, Rheology of Fibrin Clots. VI. Stress Relaxation, Creep, and Differential Dynamic Modulus of Fine Clots in Large Shearing Deformations, Journal of Rheology. 27 (1983) 135–153. https://doi.org/10.1122/1.549722.

[13] G.W. Nelb, C. Gerth, J.D. Ferry, RHEOLOGY OF FiBRIN CLOTS. III Shear creep and creep recovery of fine ligated and coarse unligated clots, Biophysical Chemistry. 5 (1976) 377–387.

[14] G.W. Nelb, G.W. Kamykowski, J.D. Ferry, Rheology of fibrin clots. v. shear modulus, creep, and creep recovery of fine unligated clots, Biophysical Chemistry. 13 (1981) 15–23. https://doi.org/10.1016/0301-4622(81)80020-8.

[15] A. Zhmurov, A.E.X. Brown, R.I. Litvinov, R.I. Dima, J.W. Weisel, V. Barsegov, Mechanism of Fibrin(ogen) Forced Unfolding, Structure. 19 (2011) 1615–1624. https://doi.org/10.1016/j.str.2011.08.013.

[16] A. Zhmurov, O. Kononova, R.I. Litvinov, R.I. Dima, V. Barsegov, J.W. Weisel, Mechanical Transition from α-Helical Coiled Coils to β-Sheets in Fibrin(ogen), J. Am. Chem. Soc. 134 (2012) 20396–20402. https://doi.org/10.1021/ja3076428.

[17] A. Zhmurov, A.D. Protopopova, R.I. Litvinov, P. Zhukov, A.R. Mukhitov, J.W. Weisel, V. Barsegov, Structural Basis of Interfacial Flexibility in Fibrin Oligomers, Structure. 24 (2016) 1907–1917. https://doi.org/10.1016/j.str.2016.08.009.

[18] R.I. Litvinov, D.A. Faizullin, Y.F. Zuev, J.W. Weisel, The α-Helix to β-Sheet Transition in Stretched and Compressed Hydrated Fibrin Clots, Biophysical Journal. 103 (2012) 1020–1027. https://doi.org/10.1016/j.bpj.2012.07.046.

[19] N.A. Kurniawan, B.E. Vos, A. Biebricher, G.J.L. Wuite, E.J.G. Peterman, G.H. Koenderink, Fibrin Networks Support Recurring Mechanical Loads by Adapting their Structure across Multiple Scales, Biophysical Journal. 111 (2016) 1026–1034. https://doi.org/10.1016/j.bpj.2016.06.034.

[20] B.E. Vos, L.C. Liebrand, M. Vahabi, A. Biebricher, G.J.L. Wuite, E.J.G. Peterman, N.A. Kurniawan, F.C. MacKintosh, G.H. Koenderink, Programming the mechanics of cohesive fiber networks by compression, Soft Matter. 13 (2017) 8886–8893. https://doi.org/10.1039/C7SM01393K.

[21] B.E. Vos, C. Martinez-Torres, F. Burla, J.W. Weisel, G.H. Koenderink, Revealing the molecular origins of fibrin’s elastomeric properties by in situ X-ray scattering, Acta Biomaterialia. 104 (2020) 39–52. https://doi.org/10.1016/j.actbio.2020.01.002.

[22] S. Munster, L.M. Jawerth, B.A. Leslie, J.I. Weitz, B. Fabry, D.A. Weitz, Strain history dependence of the nonlinear stress response of fibrin and collagen networks, Proceedings of the National Academy of Sciences. 110 (2013) 12197–12202. https://doi.org/10.1073/pnas.1222787110.

[23] F. Fleissner, M. Bonn, S.H. Parekh, Microscale spatial heterogeneity of protein structural transitions in fibrin matrices, Sci. Adv. 2 (2016) e1501778. https://doi.org/10.1126/sciadv.1501778.

[24] A.E.X. Brown, R.I. Litvinov, D.E. Discher, P.K. Purohit, J.W. Weisel, Multiscale Mechanics of Fibrin Polymer: Gel Stretching with Protein Unfolding and Loss of Water, Science. 325 (2009) 741–744. https://doi.org/10.1126/science.1172484.

[25] N.A. Kurniawan, J. Grimbergen, J. Koopman, G.H. Koenderink, Factor XIII stiffens fibrin clots by causing fiber compaction, J Thromb Haemost. 12 (2014) 1687–1696. https://doi.org/10.1111/jth.12705.

[26] J.-P. Collet, H. Shuman, R.E. Ledger, S. Lee, J.W. Weisel, The elasticity of an individual fibrin fiber in a clot, Proceedings of the National Academy of Sciences. 102 (2005) 9133–9137. https://doi.org/10.1073/pnas.0504120102.

[27] K.F. Standeven, A.M. Carter, P.J. Grant, J.W. Weisel, I. Chernysh, L. Masova, S.T. Lord, R.A.S. Ariëns, Functional analysis of fibrin γ-chain cross-linking by activated factor XIII: determination of a cross-linking pattern that maximizes clot stiffness, Blood. 110 (2007) 902–907. https://doi.org/10.1182/blood-2007-01-066837.

[28] C.C. Helms, R.A.S. Ariëns, S. Uitte de Willige, K.F. Standeven, M. Guthold, α−α Cross-Links Increase Fibrin Fiber Elasticity and Stiffness, Biophysical Journal. 102 (2012) 168–175. https://doi.org/10.1016/j.bpj.2011.11.4016.

[29] B.B.C. Lim, E.H. Lee, M. Sotomayor, K. Schulten, Molecular Basis of Fibrin Clot Elasticity, Structure. 16 (2008) 449–459. https://doi.org/10.1016/j.str.2007.12.019.

[30] N.E. Hudson, J.R. Houser, E.T. O’Brien, R.M. Taylor, R. Superfine, S.T. Lord, M.R. Falvo, Stiffening of Individual Fibrin Fibers Equitably Distributes Strain and Strengthens Networks, Biophysical Journal. 98 (2010) 1632–1640. https://doi.org/10.1016/j.bpj.2009.12.4312.

[31] J.R. Houser, N.E. Hudson, L. Ping, E.T. O’Brien, R. Superfine, S.T. Lord, M.R. Falvo, Evidence that αC Region Is Origin of Low Modulus, High Extensibility, and Strain Stiffening in Fibrin Fibers, Biophysical Journal. 99 (2010) 3038–3047. https://doi.org/10.1016/j.bpj.2010.08.060.

[32] R.D. Averett, B. Menn, E.H. Lee, C.C. Helms, T. Barker, M. Guthold, A Modular Fibrinogen Model that Captures the Stress-Strain Behavior of Fibrin Fibers, Biophysical Journal. 103 (2012) 1537–1544. https://doi.org/10.1016/j.bpj.2012.08.038.

[33] W. Liu, Fibrin Fibers Have Extraordinary Extensibility and Elasticity, Science. 313 (2006) 634–634. https://doi.org/10.1126/science.1127317.

[34] W. Liu, C.R. Carlisle, E.A. Sparks, M. Guthold, The mechanical properties of single fibrin fibers, Journal of Thrombosis and Haemostasis. (2010). https://doi.org/10.1111/j.1538-7836.2010.03745.x.

[35] M. Guthold, W. Liu, E.A. Sparks, L.M. Jawerth, L. Peng, M. Falvo, R. Superfine, R.R. Hantgan, S.T. Lord, A Comparison of the Mechanical and Structural Properties of Fibrin Fibers with Other Protein Fibers, Cell Biochem Biophys. 49 (2007) 165–181. https://doi.org/10.1007/s12013-007-9001-4.

[36] Y. Veklich, C.W. Francis, J. White, J.W. Weisel, Structural Studies of Fibrinolysis by Electron Microscopy, Blood. 92 (1998) 4721–4729.

[37] Z. Püspöki, M. Storath, D. Sage, M. Unser, Transforms and Operators for Directional Bioimage Analysis: A Survey, in: W.H. De Vos, S. Munck, J.-P. Timmermans (Eds.), Focus on Bio-Image Informatics, Springer International Publishing, 2016: pp. 69–93.

[38] R. Rezakhaniha, A. Agianniotis, J.T.C. Schrauwen, A. Griffa, D. Sage, C.V.C. Bouten, F.N. van de Vosse, M. Unser, N. Stergiopulos, Experimental Investigation of Collagen Waviness and Orientation in the Arterial Adventitia Using Confocal Laser Scanning Microscopy, Biomechanics and Modeling in Mechanobiology. 11 (2012) 461–473.

[39] A. Zumbusch, G.R. Holtom, X.S. Xie, Three-Dimensional Vibrational Imaging by Coherent Anti-Stokes Raman Scattering, Phys. Rev. Lett. 82 (1999) 4142–4145. https://doi.org/10.1103/PhysRevLett.82.4142.

[40] S.H. Parekh, Y.J. Lee, K.A. Aamer, M.T. Cicerone, Label-Free Cellular Imaging by Broadband Coherent Anti-Stokes Raman Scattering Microscopy, Biophysical Journal. 99 (2010) 2695–2704. https://doi.org/10.1016/j.bpj.2010.08.009.

[41] J.P.R. Day, G. Rago, K.F. Domke, K.P. Velikov, M. Bonn, Label-Free Imaging of Lipophilic Bioactive Molecules during Lipid Digestion by Multiplex Coherent Anti-Stokes Raman Scattering Microspectroscopy, J. Am. Chem. Soc. 132 (2010) 8433–8439. https://doi.org/10.1021/ja102069d.

[42] A.C.S. Talari, Z. Movasaghi, S. Rehman, I. ur Rehman, Raman Spectroscopy of Biological Tissues, Applied Spectroscopy Reviews. 50 (2015) 46–111. https://doi.org/10.1080/05704928.2014.923902.

[43] N.K. Howell, G. Arteaga, S. Nakai, E.C.Y. Li-Chan, Raman Spectral Analysis in the C−H Stretching Region of Proteins and Amino Acids for Investigation of Hydrophobic Interactions, J. Agric. Food Chem. 47 (1999) 924–933. https://doi.org/10.1021/jf981074l.

[44] H.G.M. Edwards, D.E. Hunt, M.G. Sibley, FT-Raman spectroscopic study of keratotic materials: horn, hoof and tortoiseshell, Spectrochimica Acta Part A: Molecular and Biomolecular Spectroscopy. 54 (1998) 745–757. https://doi.org/10.1016/S1386-1425(98)00013-4.

[45] A. Synytsya, P. Alexa, J. Besserer, J. De Boer, S. Froschauer, R. Gerlach, M. Loewe, M. Moosburger, I. Obstová, P. Quicken, B. Sosna, K. Volka, M. Würkner, Raman spectroscopy of tissue samples irradiated by protons, International Journal of Radiation Biology. 80 (2004) 581–591. https://doi.org/10.1080/09553000412331283515.

[46] S.P. Verma, D.F.H. Wallach, Raman spectra of some saturated, unsaturated and deuterated C18 fatty acids in the HCH-deformation and CH-stretching regions, Biochimica et Biophysica Acta (BBA) - Lipids and Lipid Metabolism. 486 (1977) 217–227. https://doi.org/10.1016/0005-2760(77)90018-2.

[47] Z. Movasaghi, S. Rehman, I.U. Rehman, Raman Spectroscopy of Biological Tissues, Applied Spectroscopy Reviews. 42 (2007) 493–541. https://doi.org/10.1080/05704920701551530.

[48] M. Berjot, J. Marx, A.J.P. Alix, Determination of the secondary structure of proteins from the Raman amide I band: The reference intensity profiles method, J. Raman Spectrosc. 18 (1987) 289–300. https://doi.org/10.1002/jrs.1250180411.

[49] B. Hernández, F. Pflüger, S.G. Kruglik, M. Ghomi, Characteristic Raman lines of phenylalanine analyzed by a multiconformational approach: Multiconformational analysis of the zwitterionic Phe Raman data, J. Raman Spectrosc. 44 (2013) 827–833. https://doi.org/10.1002/jrs.4290.

[50] P.R. Onck, T. Koeman, T. van Dillen, E. van der Giessen, Alternative explanation of stiffening in cross-linked semiflexible networks, Phys. Rev. Lett. 95 (2005) 178102. https://doi.org/10.1103/PhysRevLett.95.178102.

[51] B. Hernández, Y.-M. Coïc, F. Pflüger, S.G. Kruglik, M. Ghomi, All characteristic Raman markers of tyrosine and tyrosinate originate from phenol ring fundamental vibrations: Characteristic Raman markers of tyrosine and tyrosinate, J. Raman Spectrosc. 47 (2016) 210–220. https://doi.org/10.1002/jrs.4776.

[52] G. Sieler, R. Schweitzer-Stenner, The Amide I Mode of Peptides in Aqueous Solution Involves Vibrational Coupling between the Peptide Group and Water Molecules of the Hydration Shell, J. Am. Chem. Soc. 119 (1997) 1720–1726. https://doi.org/10.1021/ja960889c.

[53] M. Biancalana, S. Koide, Molecular mechanism of Thioflavin-T binding to amyloid fibrils, Biochimica et Biophysica Acta (BBA) - Proteins and Proteomics. 1804 (2010) 1405–1412. https://doi.org/10.1016/j.bbapap.2010.04.001.

[54] A.I. Sulatskaya, A.V. Lavysh, A.A. Maskevich, I.M. Kuznetsova, K.K. Turoverov, Thioflavin T fluoresces as excimer in highly concentrated aqueous solutions and as monomer being incorporated in amyloid fibrils, Sci Rep. 7 (2017) 2146. https://doi.org/10.1038/s41598-017-02237-7.

[55] C. Xue, T.Y. Lin, D. Chang, Z. Guo, Thioflavin T as an amyloid dye: fibril quantification, optimal concentration and effect on aggregation, R Soc Open Sci. 4 (2017) 160696. https://doi.org/10.1098/rsos.160696.

[56] F. Wei, D. Zhang, N.J. Halas, J.D. Hartgerink, Aromatic Amino Acids Providing Characteristic Motifs in the Raman and SERS Spectroscopy of Peptides, J. Phys. Chem. B. 112 (2008) 9158–9164. https://doi.org/10.1021/jp8025732.

[57] R.F. Doolittle, Structural basis of the fibrinogen–fibrin transformation: contributions from X-ray crystallography, Blood Reviews. 17 (2003) 33–41. https://doi.org/10.1016/S0268-960X(02)00060-7.

[58] R.F. Doolittle, X-ray crystallographic studies on fibrinogen and fibrin, J Thromb Haemost. 1 (2003) 1559–1565. https://doi.org/10.1046/j.1538-7836.2003.00278.x.

[59] J.M. Kollman, L. Pandi, M.R. Sawaya, M. Riley, R.F. Doolittle, Crystal Structure of Human Fibrinogen, Biochemistry. 48 (2009) 3877–3886. https://doi.org/10.1021/bi802205g.

[60] FGA - Fibrinogen alpha chain precursor - Homo sapiens (Human) - FGA gene & protein, (n.d.). https://www.uniprot.org/uniprot/P02671 (accessed March 28, 2020).

[61] FGB - Fibrinogen beta chain precursor - Homo sapiens (Human) - FGB gene & protein, (n.d.). https://www.uniprot.org/uniprot/P02675 (accessed March 28, 2020).

[62] FGG - Fibrinogen gamma chain precursor - Homo sapiens (Human) - FGG gene & protein, (n.d.). https://www.uniprot.org/uniprot/P02679 (accessed March 28, 2020).

[63] E. Da Silva, D. Rousseau, Molecular order and thermodynamics of the solid–liquid transition in triglycerides via Raman spectroscopy, Phys. Chem. Chem. Phys. 10 (2008) 4606. https://doi.org/10.1039/b717412h.

[64] E. Da Silva, S. Bresson, D. Rousseau, Characterization of the three major polymorphic forms and liquid state of tristearin by Raman spectroscopy, Chemistry and Physics of Lipids. 157 (2009) 113–119. https://doi.org/10.1016/j.chemphyslip.2008.11.002.

[65] C.J. Orendorff, M.W. Ducey, J.E. Pemberton, Quantitative Correlation of Raman Spectral Indicators in Determining Conformational Order in Alkyl Chains, J. Phys. Chem. A. 106 (2002) 6991–6998. https://doi.org/10.1021/jp014311n.

[66] Y. Takahashi, L. Puppulin, W. Zhu, G. Pezzotti, Raman tensor analysis of ultra-high molecular weight polyethylene and its application to study retrieved hip joint components, Acta Biomaterialia. 6 (2010) 3583–3594. https://doi.org/10.1016/j.actbio.2010.02.051.

[67] H. Wu, J.V. Volponi, A.E. Oliver, A.N. Parikh, B.A. Simmons, S. Singh, In vivo lipidomics using single-cell Raman spectroscopy, Proc Natl Acad Sci U S A. 108 (2011) 3809–3814. https://doi.org/10.1073/pnas.1009043108.

[68] M. Motoyama, M. Ando, K. Sasaki, H.-O. Hamaguchi, Differentiation of Animal Fats from Different Origins: Use of Polymorphic Features Detected by Raman Spectroscopy, Appl Spectrosc. 64 (2010) 1244–1250. https://doi.org/10.1366/000370210793335070.

[69] M. Motoyama, K. Chikuni, T. Narita, K. Aikawa, K. Sasaki, In Situ Raman Spectrometric Analysis of Crystallinity and Crystal Polymorphism of Fat in Porcine Adipose Tissue, J. Agric. Food Chem. 61 (2013) 69–75. https://doi.org/10.1021/jf3034896.

[70] M. Motoyama, I. Nakajima, H. Ohmori, G. Watanabe, K. Sasaki, A Raman Spectroscopic Method of Evaluating Fat Crystalline States and Its Application in Detecting Pork Fat, JARQ. 52 (2018) 17–22. https://doi.org/10.6090/jarq.52.17.

