## Supplementary Informatio for "Effect of shear and tensile loading on fibrin molecular structure revealed by coherent Raman microscopy"

Table of contents of SI information

- Fig. S1: The geometry optics of the calculating heights.
- Equation S1 and S2 for geometry optics of calculating heights.
- Figure S2: Fibrin constructs for microscopy of shear and tensile loaded samples.
- Figure S3: Sample stages construct for mechanical test design.
- Figure S4, The force axis angle versus shear strains.
- SI Table 1, the histogram curve fit of shear deformation.
- SI Table 2, the histogram curve fit of tensile deformation.
- Equation S3, S4 and S5 for the ratio.

Supplementary Figures

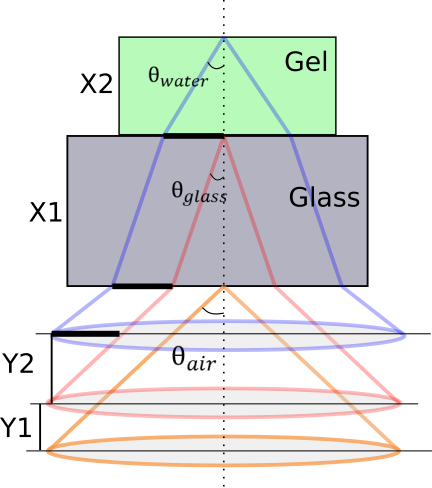

**Figure S1. Geometrical optics for calculating fibrin hydrogel height from microscopy.** Distances are described in the text below.

To obtain the height of the gel *X2*, the following geometrical relations were used. The distances *Y1* (the amount of objective displacement from the bottom of the coverslip to the fibrin/glass interface) and *Y2* (the amount of objective displacement from the bottom to the top of the fibrin hydrogel) are known from the amount the objective was translated during focusing. The thickness of the glass coverslip (*X1*) is also known. The objective for shear measurements had an NA of 0.85, and we assumed that the brightfield light filled the NA entirely. Based on geometrical optics, we can write:

${NA}_{obj}=n_{air}*sin\left( \theta_{air} \right)=n_{glass}*sin\left( \theta_{glass} \right)=n_{water}*sin\left( \theta_{water} \right)$	(Eq. S1)

where NA_obj_ = 0.85, sin *θ*_air_ = 0.85, sin *θ*_glass_ = 0.56, sin *θ*_water_ = 0.64. Thus *θ*_air_ = 58º, *θ*_glass_ = 34º and *θ*_water_ = 39º.

Because the lateral displacement of the light due to refraction is the same at the two glass interfaces (marked by the thick black lines) and *Y1* and *Y2* are known, we can relate X_2_ and Y_2_ by trigonometry to obtain distance:

$x_{2}*tan\left( \theta_{water} \right)=y_{2}*tan\left( \theta_{air} \right)$	(Eq. S2)

tan *θ*_air_ ≈ 1.60, tan *θ*_water_ ≈ 0.8, thus *X2* = Y_2_ *2.

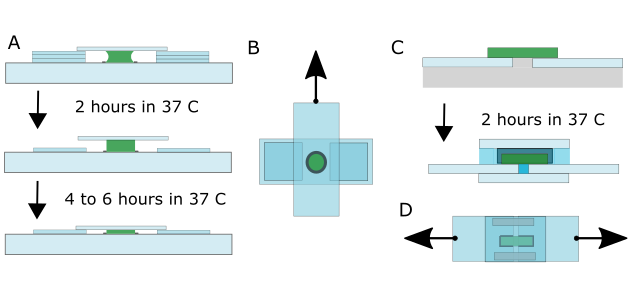
**Figure S2. Fibrin constructs for microscopy of shear and tensile loaded samples. (A)** Fibrin hydrogel preparation for shear loading. A fibrin solution was gelled between a glass slide and coverslip with three coverslips as spacers. After two hours at 37 °C, two of three coverslips were removed, and samples were left in 37 °C again for 4-6 hours. **(B)** Top view of samples for shear deformation; the top coverslip was pulled to create shear strains. **(C)** Fibrin hydrogel preparation for tensile loading. A fibrin solution was prepared across two coverslip surfaces docked to a Teflon ridge in the center. After two hours, the Teflon plate (and ridge) were removed, and the sample was sandwiched with a top and bottom coverslip and two metal spacers on either side of the gel were placed to support the top coverslip. During an experiment, we maintained humid conditions by continuous flow of buffer into the sample. **(D)** Top view of samples for tensile deformation. For actual loading stages and geometry see Figure S3.

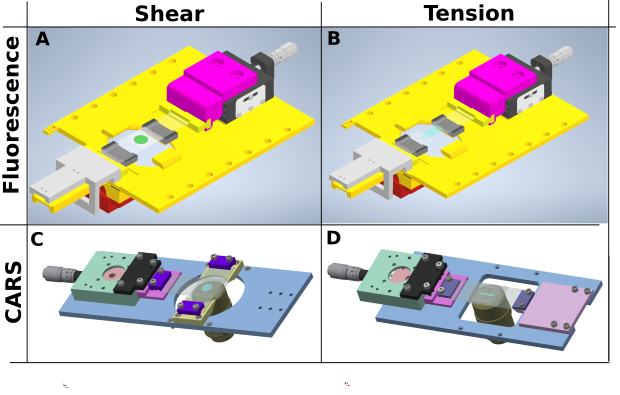
**Figure S3. Sample stages for confocal and BCARS imaging during mechanical deformation.** Two setups were used for shear and tensile loading for confocal and BCARS imaging. The colors used here are only for demonstration purposes. (**A**) Sample stage design for confocal fluorescence imaging of sheared fibrin; glass coverslips support for the fibrin sample were glued to the two (fixed) yellow stubs and to the yellow adapter attached to the translation stage. The fibrin sample (green) was sheared by moving the translation stage. (**B**) Sample stage design for confocal fluorescence imaging of tensed fibrin; glass coverslip supports for the fibrin sample were glued to the two (fixed) yellow stubs and yellow adapter attached to the translation stage. The fibrin sample (green) was stretched by moving the translation stage. In A and B, the two black plastic pieces supported the translating coverslip to maintain its height. (**C**) Sample stage design for shear deformation for BCARS; the fibrin sample sandwich was clamped down to the metal plate and translation stage by three plastic clamps (purple). The fibrin sample (blue) was stretched by moving the translation stage. (**D**) Sample stage design for tensile deformation for BCARS imaging; the fibrin sample sandwich was clamped by two plastic clamps (purple). The fibrin sample (blue) was stretched by moving the translation stage.

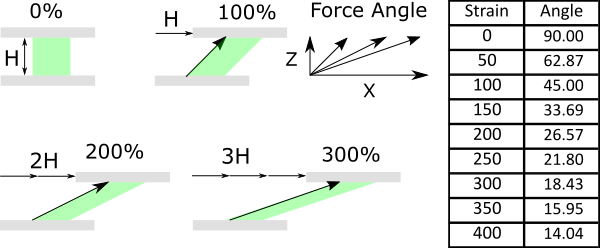

**Figure S4. The loading axis rotates in linear shear.** Shear force direction changes with increasing shear strain. (*Left*) The force angle rotates from 0%, 100%, 200% and 300% strain, and (Right) shows was a table for the loading angle (degrees) at different strain.

| **Shear** | Phenylalanine ratio | | CH_3_ ratio | | Amide I ratio | |
| --- | --- | --- | --- | --- | --- | --- |
|  | Peak position | FWHM | Peak position | FWHM | Peak position | FWHM |
| 0% | 0.1464 | 0.0627 | 0.7578 | 0.0145 | 1.2111 | 0.5015 |
| 50% | 0.1426 | 0.0665 | 0.7451 | 0.0173 | 1.3087 | 0.5182 |
| 100% | 0.1667 | 0.0636 | 0.7533 | 0.0186 | 1.1451 | 0.4568 |
| 150% | 0.1487 | 0.0714 | 0.7754 | 0.0202 | 0.9753 | 0.4254 |
| 250% | 0.2040 | 0.0898 | 0.8732 | 0.0292 | 1.1138 | 0.4009 |
| 300% | 0.2052 | 0.1022 | 0.9373 | 0.0223 | 1.1542 | 0.4034 |
| 350% | 0.2269 | 0.1050 | 0.9496 | 0.0302 | 0.9600 | 0.3526 |
| 400% | 0.2328 | 0.1197 | 0.9903 | 0.0334 | 1.3215 | 0.5063 |
| Average | 0.184 | 0.085 | 0.848 | 0.023 | 1.149 | 0.408 |

**Table S1. Curve fitting of histograms from fibrin shear deformation.** Peak position, full width at maximum (FWHM), and average value of histograms of the Phenylalanine ratio, CH_3_ ratio, and Amide I ratio for data shown in **Figure 3** of the main text. Histogram parameters were quantified by fitting histograms with Gaussians, and peak position and FWHM were reported from the fit.

| **Tension** | Phenylalanine ratio | | CH_3_ ratio | | Amide I ratio | |
| --- | --- | --- | --- | --- | --- | --- |
|  | Peak position | FWHM | Peak position | FWHM | Peak position | FWHM |
| 0% | 1.175 | 0.610 | 1.853 | 0.077 | 0.959 | 0.377 |
| 20% | 1.102 | 0.542 | 1.933 | 0.062 | 0.775 | 0.152 |
| 40% | 1.428 | 0.807 | 2.015 | 0.084 | 0.828 | 0.166 |
| 60% | 1.329 | 0.846 | 2.012 | 0.094 | 0.783 | 0.126 |
| 80% | 1.245 | 0.849 | 1.987 | 0.087 | 0.763 | 0.116 |
| 100% | 0.860 | 0.784 | 2.004 | 0.241 | 0.674 | 0.118 |
| Average | 1.190 | 0.740 | 1.967 | 0.107 | 0.797 | 0.176 |

**Table S2. Curve fitting of histograms from fibrin tensile deformation.** Peak position, (FWHM), and average value of histograms of the Phenylalanine ratio, CH_3_ ratio, and Amide I ratio for data shown in **Figure 4**. Histogram parameters were quantified by fitting with Gaussians, and peak position and FWHM were reported from the fit.

$PhenylalanineRatio=\frac{Intensityof980-1010{cm}^{-1}}{Intensityof1010-1060{cm}^{-1}}$	(Eq. S3)

$AmideIRatio=\frac{Intensityof1643-1653{cm}^{-1}}{Intensityof1669-1679{cm}^{-1}}$	(Eq. S4)

${CH}_{3}Ratio=\frac{Intensityof2920-2950{cm}^{-1}}{Intensityof2950-2980{cm}^{-1}}$ (Eq. S5)

**Equation S3, S4, and S5 for ratio from BCARS data.** The equations for Phenyl ring, Amide I and CH_3_ ratios. Intensities were integrated from the assigned Raman shifts.
